# Young bone marrow-derived microglia-like cells improve cognitive function in the aged brain through BDNF-TrkB signalling

**DOI:** 10.64898/2026.10.15.759429

**Authors:** Lukasz Wlodarek, Ren-Ke Li

## Abstract

Blood-borne macrophages are present in the brain under normal and pathological conditions. Although the complete role of these cells has yet to be determined, emerging evidence shows these cells to be vital during injury and infection, and contribute to the brain’s macrophage niche throughout the entirety of an organism’s lifespan. This is not always beneficial as macrophages become pro-inflammatory, primed, and hyper-activated with age and disease. One solution is to introduce macrophage precursor cells from young donors into the aged brain to attenuate age- and injury-induced inflammation and cognitive decline. Despite drastic improvements, it remains unclear by which mechanism(s) these cells rejuvenate brain function. To understand this, we generated mice whose CX3CR1-expressing macrophages are selectively depletable using diphtheria toxin. Controlled ablation of donor bone marrow (BM)- derived macrophages from the aged brain was achieved one or three months after transplanting into aged wild type recipients. At three months, the overall brain microglial-niche failed to repopulate from either peripherally- or centrally-derived macrophages after depletion. This was accompanied by severely reduced global monocyte recruitment and a large influx of neutrophils to the brain. Hippocampal-dependent behaviour was unaffected, presumably due to the absence of these cells in the hippocampus during this time point. At one month, BM-derived microglia-like cells were abundant in the hippocampus and their depletion severely impaired improvements in cognitive function and inhibited dendritic spine arborisation in hippocampal neurons. Pharmacological blocking of the brain-derived neurotrophic factor (BDNF) receptor, tropomyosin-related kinase B (TrkB), yielded the same detrimental effects in these mice as depleting the cells. These findings suggest that young BM-derived microglia-like cells could improve cognitive function in aged animals via the BDNF-TrkB signalling pathway during early brain homing and do not require permanent integration for beneficial effects to persist long-term.

## Introduction

Microglia are the central nervous system’s resident macrophages that act as the initial and primary form of immune defense against invading pathogens and disease [1]. Additionally, microglia have been implicated in tissue repair via the secretion of anti-inflammatory cytokines and chemokines, as well as maintaining synaptic integrity and shaping neuronal circuitry [2–4]. However with age, microglia begin to acquire a pro-inflammatory phenotype that manifests as numerous different neurobehavioral complications, impaired neuroprotection, and detrimentally exaggerated responses to injury and disease [5–7]. As such, therapies targeting aged and dysfunctional microglia are attractive avenues of treatment.

Previously we demonstrated the possibility of introducing anti-inflammatory peripherally-derived microglial-like cells to the aged brain following bone marrow (BM) reconstitution with young BM stem cell antigen-1 (Sca-1)-expressing cells [8]. We and others found that these young-to-old chimeras demonstrated improved cognitive performance, wound healing, neuroprotection, and enhanced synaptic connectivity and plasticity when compared to aged mice reconstituted with old BM cells under normal and pathological conditions [8–10]. Several different studies have identified microglial paracrine signalling to be one of the main contributors responsible for improving global aged brain function following either transplantation with young BM cells or parabiosis with a young donor [10–12]. However the question remains whether the therapeutic potential of young blood-borne microglial-like cells in the aged brain requires permanent or transient integration.

The past decade saw the emergence of techniques capable of selectively and globally depleting microglia from the brain using mice expressing tamoxifen-inducible Cre recombinase (CreER) under the control of the microglial CX_3_CR1 promoter (CX_3_CR1^CreER^) to drive diphtheria toxin receptor expression and subsequent depletion following diphtheria toxin administration [12–14], and exogenous administration of CSF1R inhibitors [15–17], respectively. Depletion of microglia from the brain is either both severely detrimental to cognitive function and repair after injury, or substantially improves cognitive performance and promotes brain recovery [18]. These polarizing results most likely stem from the fact that the age of the animal is not always taken into consideration. This is important because older animals whose microglia have been depleted seem to benefit from depletion (i.e. improved recovery after stroke, amelioration of radiation-induced cognitive deficits, reduced neuronal and memory loss in Alzheimer’s) [19–21]. In contrast, depletion of microglia from younger animals was seen to increase infarct size and brain injury after stroke, and reduce synaptic plasticity associated to learning and behavioural performance [22–24].

These studies have primarily focused on depleting brain-resident macrophages to examine the role that these cells play on neuronal function, cognition, and response to injury. Little to no attention has been paid in understanding the contribution of peripherally-derived macrophages on brain physiology and behaviour. Indeed, blood-borne macrophages have been extensively documented as having a pivotal role in wound healing, preservation of neural synapses, and preconditioning the brain for better response and recovery to insult [25–27]. In this study, we aim to investigate whether depleting young BM-derived microglia-like cells reverses the beneficial effects the cells bring to the aged and irradiated brain. This is important to elucidate as it would provide valuable insight into the parameters required to translate this microglial replacement therapy for attenuating brain dysfunction in the elderly and in patients with neurodegenerative disorders.

## Methods

### Animals

Young (2-3-month-old) C57BL/6-Tg(CAG-EGFP)1Osb/J, C57BL/6 WT, B6.129P2(Cg)-Cx3cr1tm2.1(cre/ERT2)Litt/WganJ, and C57BL/6-Gt(ROSA)26Sortm1(HBEGF)Awai/J mice were obtained from Jackson Laboratories. Mice were bred and aged to 2-20 months within the animal housing facility. All mice were kept on a 12-hour light/dark cycle and provided with free access to both food and water. The Animal Care Committee of the University Health Network approved all experimental procedures, which were carried out according to the Guide for the Care and Use of Laboratory Animals (National Institutes of Health, 2011).

### Donor cell preparation

Young and old (18-20-month-old) BM was obtained from the femurs and tibiae of C57BL/6-Tg-GFP mice and from *Cx3cr1*^CreER/+^R26^iDTR/+^ mice only young BM was obtained. The bones were flushed with sterile PBS and then incubated in RBC lysis buffer (154.42 mM NH4Cl, 11.9 mM NaHCO3, 0.026mM EDTA) for 5 min, followed by centrifugation at 1000 rpm for 5 min. The pellet was then suspended in Iscove’s Modified Dulbecco’s Medium (Thermo Fisher Scientific) and passed through a 40 µm filter. Cells were counted and Sca-1^+^ isolated using immunomagnetic activated cell sorting according to the manufacturer’s instructions (STEMCELL Technologies).

### Bone marrow reconstitution

Female C57BL/6J mice (18-20-month-old) were irradiated at 10 Gy (Cs-137 irradiator, Gammacell 40 Exactor), after which 2×10^6^ young or old GFP^+^/Sca-1^+^ cells or 2×10^6^ young YFP^+^/Sca-1^+^ cells were injected via the tail vein, generating Y^+^-O and O^+^-O chimeras, respectively.

### In vivo treatments

Tamoxifen (Sigma-Aldrich) was suspended in corn oil (Sigma-Aldrich) at 75°C and 5 mg (200 µl) was administered subcutaneously on two days separated by 48 hours. Diphtheria toxin (Sigma-Aldrich) was injected intraperitoneally one month after tamoxifen with three daily doses of 25ng/g. ANA-12 (S7745, Selleck) was injected intraperitoneally at 0.5mg/kg for five consecutive days.

### Flow cytometry

The brains were collected from mice 12 weeks post-reconstitution and were minced, digested, and immune cells were extracted as previously described [28]. The hearts were minced and digested using collagenase type II (Worthington, 2 mg/ml) at 37°C for 30 min and filtered through a 70 µm filter. Single cell suspensions were re-suspended in FACS Buffer (PBS Ca2^+^/Mg2^+^ free +1% FBS). Cells were stained with the following primary antibodies: CD45, 103128, Biolegend; CD11b, 101208, Biolegend; Ly6C, 128026, Biolegend; Ly6G, 127614, Biolegend; F4/80, 123114, Biolegend; CX3CR1, 149006, Biolegend. All primary antibody incubations were carried out for 1 h at 4°C in the dark after which cells were washed and events collected on a LSRII flow cytometer equipped with a violet laser (25 mW), blue laser (50 mW), yellow laser (100 mW), and red laser (40 mW). For identification of mouse cells, monocytes were CD45^+^/CD11b^+^/F4/80^−^/Ly6G^-^/Ly6C^hi/neg^, neutrophils CD45^+^/CD11b^+^/F4/80^−^/Ly6G^+^, and macrophages CD45^+^/CD11b^+^/F4/80^+^/CX3CR1^+^. Gating was determined by FMOs, and all data were analyzed in FlowJo (TreeStar).

### Behavioural tests

The open field test apparatus consisted of a 38 x 60 x 60 cm chamber with grey Plexiglas walls and transparent ceiling to allow for video recording. Mice were placed in the chamber for 10 min and their ambulatory distance and rearing count was tracked and analyzed by idTracker [29]. Additional analyses were performed using a custom python script, which transformed x- and y-coordinates associated to animal behaviour into the various parameters reported in the results section. The novel object recognition task was performed as previously described [30]. In short, mice underwent three consecutive days of testing, each separated by a whole 24 hours. The first day (habituation phase) involved the animals roaming free in a novel environment for 10 min under video surveillance. The second day (familiarization phase) introduced two identical copies of the same object, while the third and final day (testing phase) introduced one copy of an object from the second day and one novel object that the mice have never seen before. For both days, the mice were allowed to freely interact with each object for a total duration of 10 min and the interaction time for each object was separately measured and recorded. Recognition memory performance was defined by calculating the discrimination index (i.e., time spend interacting with novel object versus time spent interacting with familiar object). Objects were randomized between mice to determine which would be the familiar set and which would be the novel set, and were washed with 75% ethanol between animals in order to eliminate odour. Spatial reference learning and memory was carried out as described previously [31]. Briefly, the Barnes maze consisted of an elevator platform (30 cm above ground) with 20 equidistance holes and spatial cues around the circumference. Each mouse was given 90 s to find the correct hole leading to a dark chamber. The total time taken and the number of errors made before finding the correct hole was measured. Mice ran the maze under light stimulation for a total of four trials/day for five consecutive days with at least 20 minutes in between each trial. The maze was cleaned with 75% ethanol between each trial and if the mice could not complete the maze within 90 s, they were gently guided to the escape hole by hand. Object location memory was performed as described [32]. In short, this behavioural test was performed similarly to novel object recognition mentioned previously; however the testing phase involved shifting the location of similar objects, as opposed to introducing a novel object. Location memory was quantified by calculating the discrimination index derived from object interaction time. Lastly, the rotarod performance and hanging limb test were performed as previously described [33, 34].

### Immunofluorescence staining

Brains were extracted one or three months post-reconstitution and fixed in 4% paraformaldehyde overnight at 4°C before preservation in increasing concentrations of 10%, 20% and 30% sucrose in PBS (w/v). Brains underwent stereological sectioning coronally at a thickness of 20 µm on a freezing-sliding microtome, and sections were stored in cryoprotective medium (OCT, Dako) at −20°C. Frozen tissue sections were blocked with 5% (w/v) donkey serum before incubation at room temperature for 2 h with primary antibodies at the following concentrations: YFP, (1:400, A21311, Life Technologies); Ly6G, (1:400, 551459, BD Biosciences); Incubation with respective Alexa 488, 568, or 647 conjugated secondary antibodies (Life Technologies) was carried out at room temperature for 1 h with light protection. Cell nuclei were identified with 4’,6-diamidno-2-phenylindole (DAPI) at a 1:2000 for 5 min. Stained tissue slides were stored at −4°C under dark conditions.

### Cell quantification

Cell quantification was carried out on all the positive cells of interest present within a specific region of the brain (i.e., hippocampus) or within randomly selected regions of the cerebellum (1.0 mm^2^ cross-sections) and were averaged across three replicates. Fluorescent images were obtained with a Nikon Eclipse Ti fluorescent microscope using a VS120 slide scanner. Prior to analysis, images were captured at an identical exposure and set to and fixed fluorescence intensity using the Olympus cellSens imaging software. Number of cells were quantified automatically using built-in cell counting tools in the Olympus cellSens imaging software.

### Propidium iodide (PI) staining

PI staining was carried out using a 1:1000 dilution of PI (Sigma-Aldrich) that was applied to brain sections at room temperature in the dark for 20 min, followed by incubation for 5 min in 1:2000 DAPI, after which mounting media and a coverslip were applied. Stained sections were stored at 4°C in the dark and viewed using a Nikon Eclipse Ti fluorescent microscope.

### RNA extraction and RT-qPCR

Total RNA was isolated from mouse whole-brains (hippocampus) using Tri-Reagent (Sigma-Aldrich) according to the manufacturer’s instructions. Reverse transcriptase was performed using SuperScriptIII (Invitrogen) and 1 µg of total RNA served as a template for each reaction. cDNA expression was analyzed using SensiFast (Bioline) SYBR Green with the following parameters: 95°C for 2 min (95°C for 5 s; 60°C for 30 s for 40 cycles). Total RNA isolated from explanted cells from reconstituted brains was carried out using the RNAqueous-Micro Total RNA Isolation Kit (AM1931, ThermoFisher) according to the manufacturer’s instructions. RT-qPCR was performed extracted hippocampus using the RT² Profiler™ Mouse PCR Neurogenesis Array (330231, Qiagen, GeneGlobe ID - PAMM-404ZG-4) on a Roche LightCycler 480 (384-well Block). Relative expression levels were normalized to glyceraldehyde 3-phosphate dehydrogenase (GAPDH).

### Protein extraction and ELISA

Total protein was extracted from homogenized hippocampus and cells in lysis buffer. Soluble factor detection and quantification was carried out according to the manufacturer’s instructions: BDNF (MBS032748, MyBioSource); IL-10 (BMS614INST, ThermoFischer); PTN (MBS355435, MyBioSource); TNFα (BMS607-3, ThermoFisher); VEGFa (BMS619-2, ThermoFisher).

### Dendritic spine analysis

To study the quantity and morphology of dendritic spines of pyramidal cells in the CA1 region of the hippocampus, brains were removed from the animals and incubated in a series of solutions using the FD Rapid GolgiStain kit (FD Neurotechnologies), according to the manufacturer’s instructions. Following cryoprotection of the whole-brains with Solution C (see kit), tissue was sectioned coronally at 200 µm thickness and then viewed under a light microscope at 100x magnification. Dendritic arbours of each cell was analyzed as previously described [35]. The spine density was calculated as the number of spines (defined as protrusions of the dendritic membrane regardless of shape) along a 30 µm dendritic terminal segment. A total of 10 different neurons were analyzed for each animal per group.

### Statistical analysis

All values are expressed as mean ± s.e.m. Analyses were performed using GraphPad Prism 6.0 software. Student’s and two-tailed was used for two-group comparison, whereas a one-way ANOVA was used for three or more group comparisons. Statistical analysis of cell quantification and behavioural data was carried out using an unpaired, two-tailed t-test. Two-way ANOVA was carried out for two-factor variables followed by Tukey post-hoc tests.

## Results

### Differentiating blood-borne from brain-resident macrophages

The origin of microglia continues to be a highly debated topic. Currently, there seems to be a consensus that the vast majority of brain-resident microglia originate during development from the yolk-sac [36, 37]. Nonetheless, functional microglial-like cells have been shown to migrate from the blood and BM to the brain under normal and pathological conditions [8–10]. Recently, differentiating these two types of cells from one another has been made easier as it has been shown that brain-resident microglia are CD45^+^CD11b^+^CX3CR1^+^F4/80^low^, whereas peripherally-derived microglia-like cells are CD45^+^CD11b^+^CX3CR1^+^F4/80^hi^ [38]. We confirmed these findings in wild type and reconstituted chimeras that give rise to green fluorescent protein (GFP)-expressing microglial-like cells in the brain (Fig. 1). Here we found that before BM reconstitution with Sca-1^+^ cells, F4/80^low^ macrophages dominate (∼85%; Fig. 1a, b, g), whereas after reconstitution there seems to be a shift towards an equal percent of F4/80^hi^ and F4/80^low^ macrophages (Fig. 1c, d, g). Further examination of the CD45^+^CD11b^+^CX3CR1^+^GFP^+^ population reveals the significant majority being composed of F4/80^hi^ cells (Fig. 1e, h, i), as well as a minor population of lesser defined F4/80^low^ cells (Fig. 1f, h, i), as previously reported [38].

**Figure 1:**
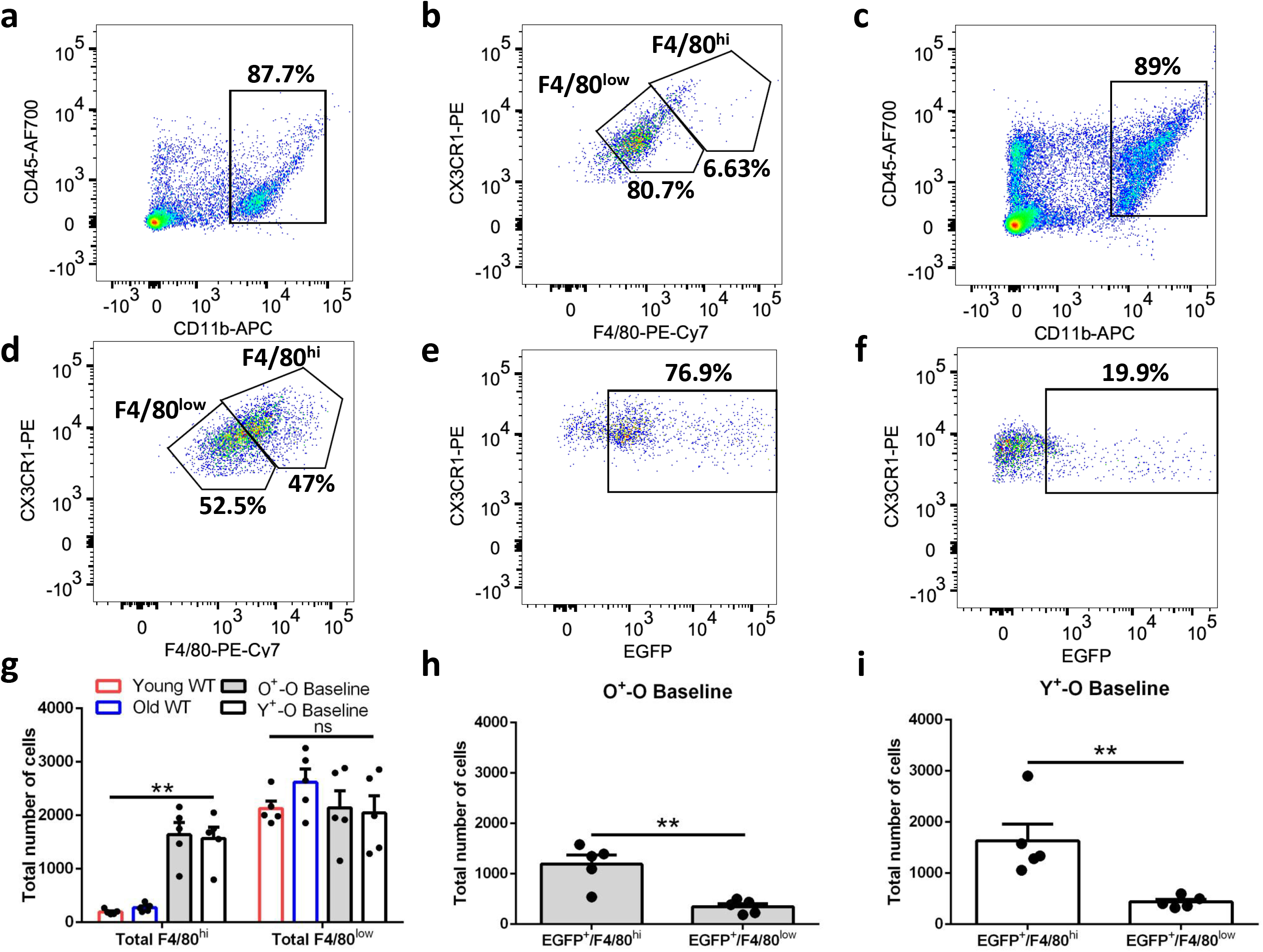
Differentiating between brain-resident and BM-derived microglia-like cells before and after transplantation with Sca-1^+^ cells using multi-color flow cytometry. (a) Total macrophage population in the brain of wild type animals organized into (b) F4/80^low^ brain-resident macrophages and F4/80^hi^ peripherally-derived macrophages. (c) Total macrophage population in reconstituted animals organized into (d) F4/80^low^ brain-resident macrophages and F4/80^hi^ peripherally-derived macrophages. Of the total number of peripherally-derived macrophages (GFP^+^), the majority of them are the expected F4/80^hi^ population (e), with a few lesser defined F4/80^low^ population (f). Quantification of cell number of total F4/80^hi^ and F4/80^low^ population (g) and further classification of F4/80^hi^ (h) and F4/80^low^ (i) population of cells that are also GFP^+^. *n* = 5 mice/group.

### Microglia depletion in *Cx3cr1*^CreER/+^R26^iDTR/+^

To achieve selective depletion of BM-derived microglial-like cells, we first bred mice with tamoxifen-inducible Cre under the microglia-expressed CX3CR1 promoter (*Cx3cr1*^CreER-EYFP^ mice) with *R26*^iDTR/+^ mice, which results in expression of diphtheria toxin receptor upon Cre recombination (Fig. 2a). Before selectively ablating BM-derived microglial-like cells from reconstituted aged recipients, we first verified depletion in *Cx3cr1*^CreER/+^R26^iDTR/+^ mice prior to BM transplantation. Upon tamoxifen and diphtheria toxin administration, we found that we were unable to significantly lower the F4/80^hi^ population away from baseline (Fig. 2b). This was expected as our previous findings demonstrated very few of these cells in the brain before reconstitution. Next we looked at whether we were able to ablate F4/80^low^ cells and found that this population declined by ∼60% from baseline 7 days after injection with diphtheria toxin (Fig. 2c). Interestingly, following depletion of CD45^+^CD11b^+^CX3CR1^+^YFP^+^F4/80^+^ cells, we noticed a significant increase in non-depletable blood-borne macrophages (YFP^-^/F4/80^hi^) in the brain relative to baseline (Fig. 2d), but not in non-depletable brain-residence macrophages (YFP^-^/F4/80^low^; Fig. 2e). Lastly, we quantified depletion of total depletable macrophages (YFP^+^/F4/80^+^) in the heart as an internal control and found a significant decline in the macrophage population by day 3 (∼75%), confirming our model, despite noticing a spike in cell numbers by day 7 (Fig. 2f). This finding is likely the result of artificially inflated YFP signal from persisting dead cells that have not been cleared, in addition to rapid expansion of the macrophage niche, as several different groups have reported impaired clearance of tissue debris and dead cells following depletion of macrophages from the heart [39].

**Figure 2:**
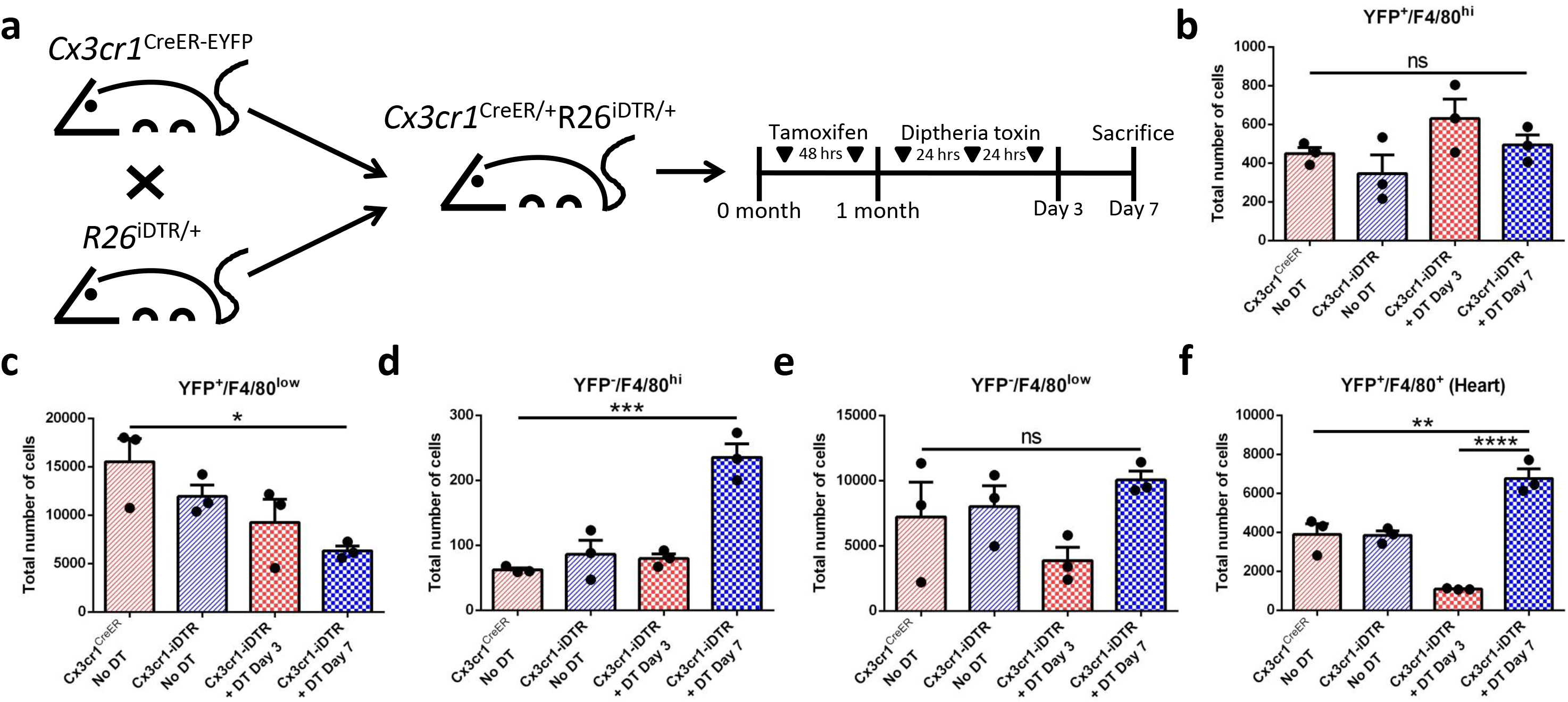
Generation of transgenic mice whose CX3CR1^+^ microglia are depletable following tamoxifen and diphtheria toxin administration. (a) Cartoon representation of mouse breeding for generation of *Cx3cr1*^CreER/+^R26^iDTR/+^ mice. (b) Depletable CX3CR1^+^ microglial cells (YFP^+^) that are F4/80^hi^ and F4/80^low^ (c) during day 3 and day 7 depletion time points in the brain. (d) Non-depletable (YFP-) microglia during both depletion time points in the brain that are F4/80^hi^ and F4/80^low^ (e). (f) Using the heart as an internal control to validate macrophage depletion. *n* = 3 mice/group.

### Selective depletion and repopulation of BM-derived microglia-like cells

After validating depletion from *Cx3cr1*^CreER/+^R26^iDTR/+^ mice, we then took the BM of these animals, purified the Sca-1 population of cells, and reconstituted aged animals with these cells from young donors, generating Y^+^-O chimeras (Fig. 3a). Three months following BM reconstitution, we were able to deplete an average of 72%, 55%, and 60% of BM-derived (YFP^+^) total, F4/80^hi^, and F4/80^low^ macrophages from the brain by day 7, respectively (Fig. 3b-d). This time point was selected based on our previous data and findings from other groups that likewise demonstrated peak depletion by this day [38, 40]. Interesting, in stark contrast to numerous previously published microglia depletion studies [41], to our knowledge we are the first group to report not witnessing repopulation of the microglial niche from the periphery or from brain-resident cells by day 28 (a repopulation time point used by most studies) in reconstituted aged animals (Fig. 3e, f). This failure to repopulate the microglial niche from the BM after depletion may stem from significantly attenuated recruitment of monocytes to the brain by day 28 (Fig. 3g). Furthermore, impaired expansion of the microglial niche after depletion may also be inhibited by a significant surge of neutrophils to the brain during the repopulation window (Fig. 3h).

**Figure 3:**
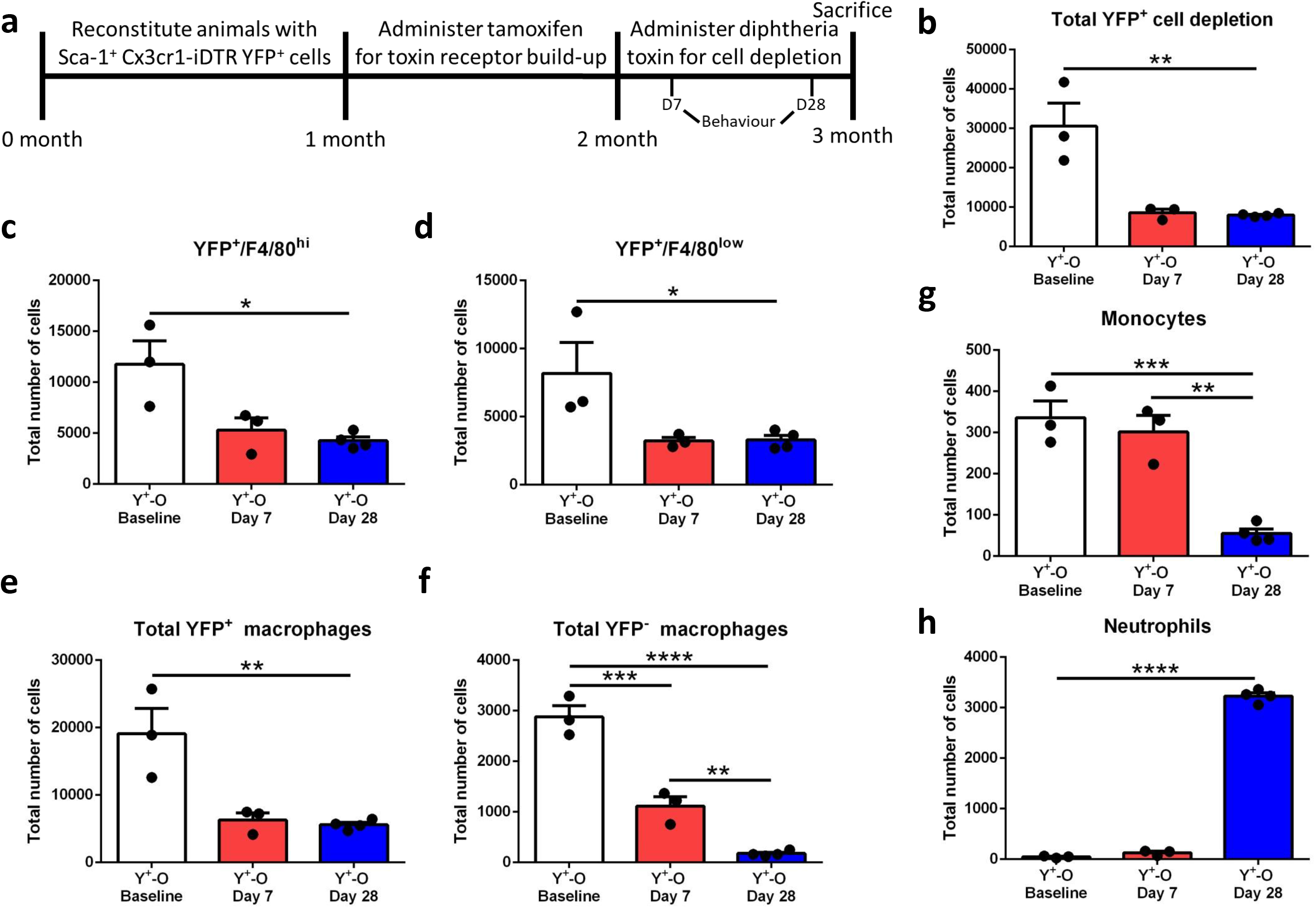
Selective ablation of young BM-derived microglia-like cells from the aged brain. (a) Timeline for BM transplantation, diphtheria toxin administration and behavior during depletion and repopulation. (b) Total BM-derived (YFP^+^) cell depletion, irrespective of phenotype, subcategorized into the F4/80^hi^ (c) and F4/80^low^ (d) population of cells. (e) Total number of YFP^+^ and YFP^-^ (f) macrophages before and after depletion. (g) Changes in the number of monocytes and neutrophils (h) in response to microglial depletion and repopulation.

### Selective depletion of BM-derived microglia-like cells alters cognitive function

Previously we found that blood-borne microglia-like cells populate the brain after BM reconstitution with Sca-1^+^ cells and aged mice that received peripherally-derived macrophages from young donors demonstrated significantly better cognitive performance, synaptic plasticity, resilience against injury, and improved tissue repair [8]. In this study, we partially depleted BM-derived macrophages (CD45^+^CD11b^+^CX3CR1^+^F4/80^hi^) 3-months post-reconstitution to determine whether short- or long-term integration of cells is required for the beneficial cognitive effects to persist. We found that depleting the cells led to a significant effect on overall activity levels (Fig. 4a, b), fear (Fig. 4c), and anxiety by day 7 and by day 28 (Fig. 4d). Despite this, performance on hippocampus-dependent behavioural tasks, such as: rearing (Fig. 4e), spatial learning and memory (Fig. 4f, g, i), and recognition memory (Fig. 4h), did not significantly decline from baseline by day 7 or by day 28. In contrast to this, grip strength (Fig. 4j) and motor performance and coordination (Fig. 4k) displayed greatest decline from baseline following partial ablation of BM-derived microglial-like cells.

**Figure 4:**
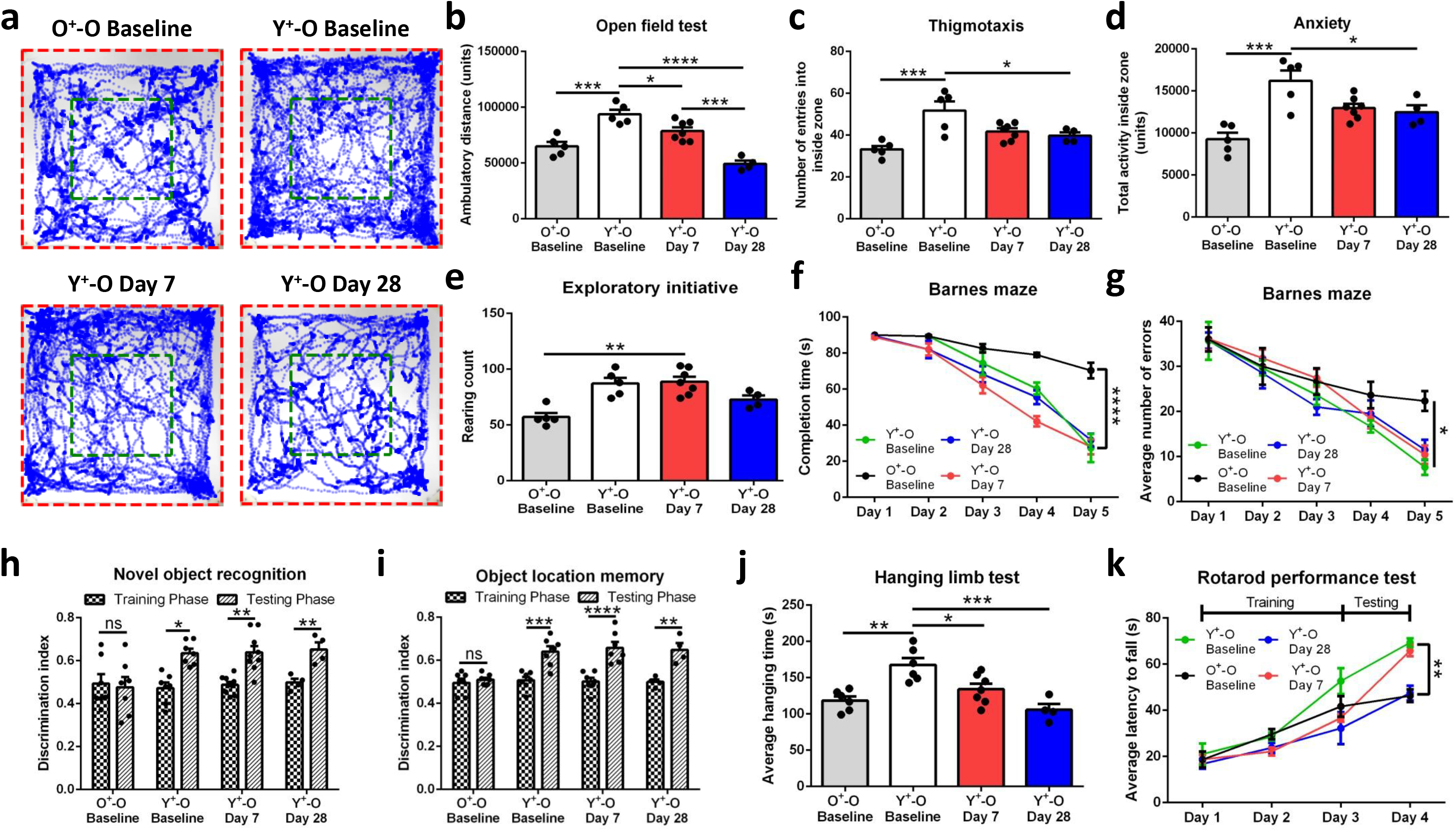
Hippocampal and cerebellar behavioral analysis following ablation of BM-derived microglia-like cells during depletion and repopulation time points. (a) Open field test analysis with representative traces and (b) quantified activity levels. (c) Measurement of movement from outside to inside zones of the measuring apparatus (number of entries) complemented by total activity within the inside zone (d). Exploratory/curiosity initiative quantified by total number of times the animal rears (e). Barnes maze spatial learning and memory behavioral paradigm measuring maze completion time (f) and average number of errors made before finding the correct escape hole (g). Novel object recognition (h) and object location spatial memory (i) quantified by a ratio of discrimination ability between novel objects or similar objects with dynamic location. (j) Hanging limb test and (k) rotarod performance test for cerebellar function. *n* = 6 mice/group.

Taken together, it remains unclear whether this decline in function is the result of: (1) selective ablation of young BM cells; (2) a decrease in the recruitment of macrophage precursor cells during repopulation; or (3) the triggering of global inflammation by day 28 following cell depletion. Nonetheless, despite previously reporting decreased recruitment of monocytes and a massive influx of neutrophils during repopulation, partial depletion of young BM-derived microglia-like cells in the aged brain did not reverse improvements in hippocampal-driven learning and memory to levels present in aged animals whose BM was reconstituted with old cells.

### BM-derived microglia-like cells persist after diphtheria toxin in the cerebellum

In order to understand why cerebellar function declined after depletion and did not return back to baseline following repopulation, we sectioned different areas of the cerebellum and found YFP^+^ cells during day 7 and day 28 with altered cell morphology when compared to baseline (Fig. 5a, c, d). Surprisingly, we did not see a decline in YFP^+^ cells following depletion, even though the cells displayed necrotic/apoptotic morphological appearance at day 7 and day 28 (Fig. 5a, c). In the hippocampus, we were unable to detect a large number of YFP^+^ cells 3-months post-reconstitution (Fig 5b, e), possibly giving a clue as to why hippocampal-dependent learning and memory does not decline at this time point following depletion.

**Figure 5:**
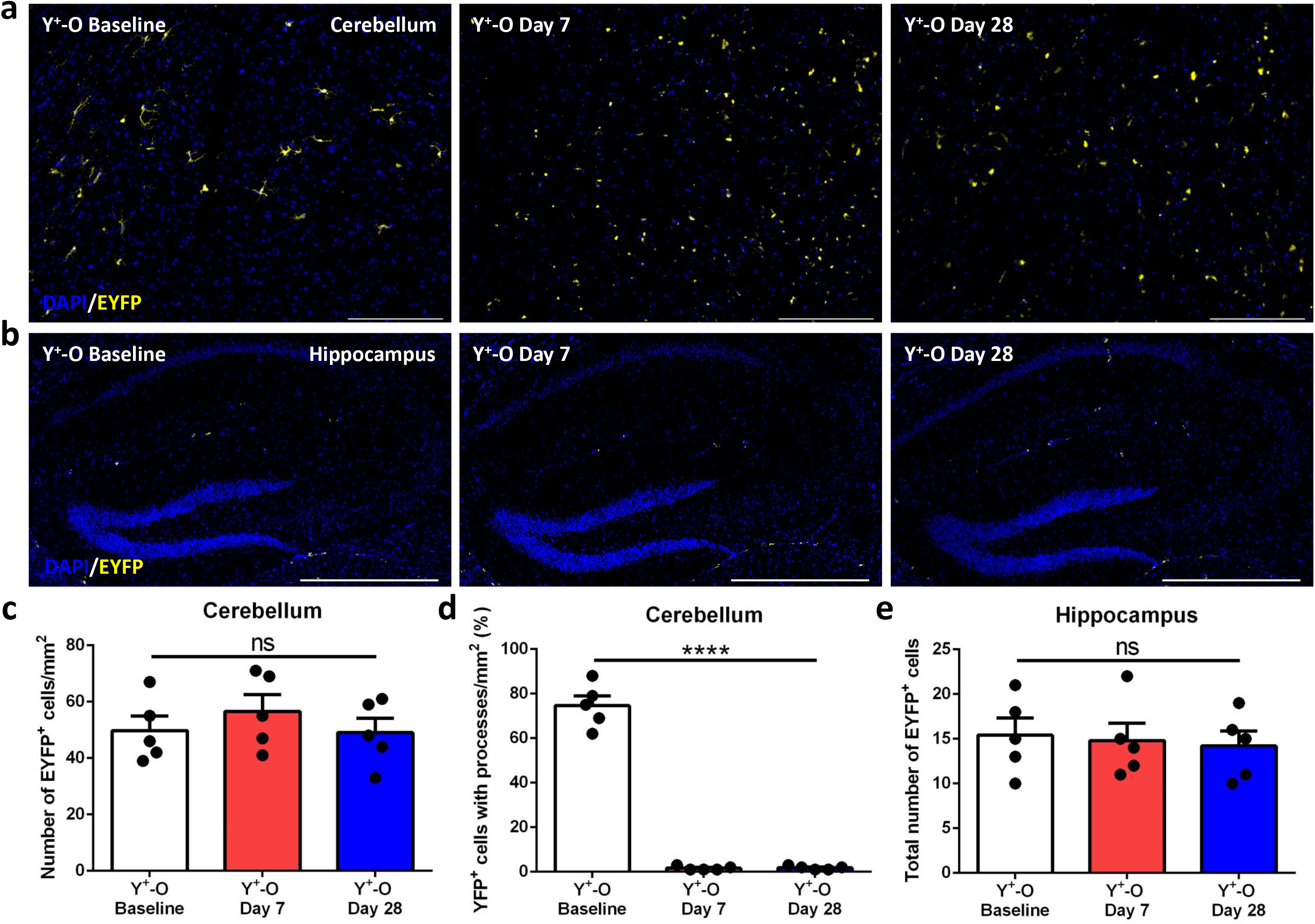
Persistence of YFP^+^ signal in the cerebellum following diphtheria toxin. (a) Immunofluorescence staining of BM-derived YFP^+^ cells in the cerebellum and (b) hippocampus during depletion and repopulation. (c) Quantification of number of YFP^+^ cells in the cerebellum and the number of cells approximating a viable cell based on processes protruding from the cell body (d). (e) Quantification of total number of YFP^+^ cells in the hippocampus. Scale bars 200 µm with *n* = 5 mice/group.

### Aged mice demonstrate impaired clearance of dead BM-derived microglia-like cells

Numerous previous studies using this model of depletion were able to achieve >80% depletion efficiency using young (∼2-month-old) mice [18]. We were curious to understand whether our lower depletion efficiency was the result of these experiments performed on 21-month-old mice. Indeed, toxin-induced cell death requires clearance for physical emptying of the microglial niche, which then allows for subsequent recruitment of monocytes and differentiation into microglia for repopulation of the CX3CR1^+^ compartment. We found that after administration of diphtheria toxin, there was a significant number of Ly6G^+^ neutrophils present within the vicinity of YFP^+^ cells during depletion and repopulation, relative to baseline (Fig. 6a-d). At day 28 it was also observed that neutrophils were frequently expressing YFP (Fig. 6c, e). Since CX3CR1 is not expressed on neutrophils, these findings suggest the occurrence of phagocytosis. Indeed, most YFP^+^ cells persisting following depletion are positive for propidium iodide (PI), a DNA-binding fluorescent dye that enters cells with a compromised cell membrane during depletion and repopulation time points (Fig. 6f-i). Taken together, these findings demonstrate that depleting BM-derived microglia-like cells from the brains of aged animals is challenging due to impaired clearance of these cells for up to nearly 1 month after depletion. The continual presence of YFP^+^PI^+^ macrophages this long after depletion may explain our much lower depletion efficiency when compared to other papers [18], which would likely be skewed by false positive counting of dead YFP^+^ cells as living due to seemingly age-dependent impairments in cell and debris clearance. Lastly, macrophage depletion in the brain is challenging due to the cell’s resilience and rapid repopulation 48-72 hours after initial depletion [42]. Our findings demonstrate failure for repopulation of this niche. A plausible explanation for this may be due to physical occupation of the microglial niche by dead cells after depletion.

**Figure 6:**
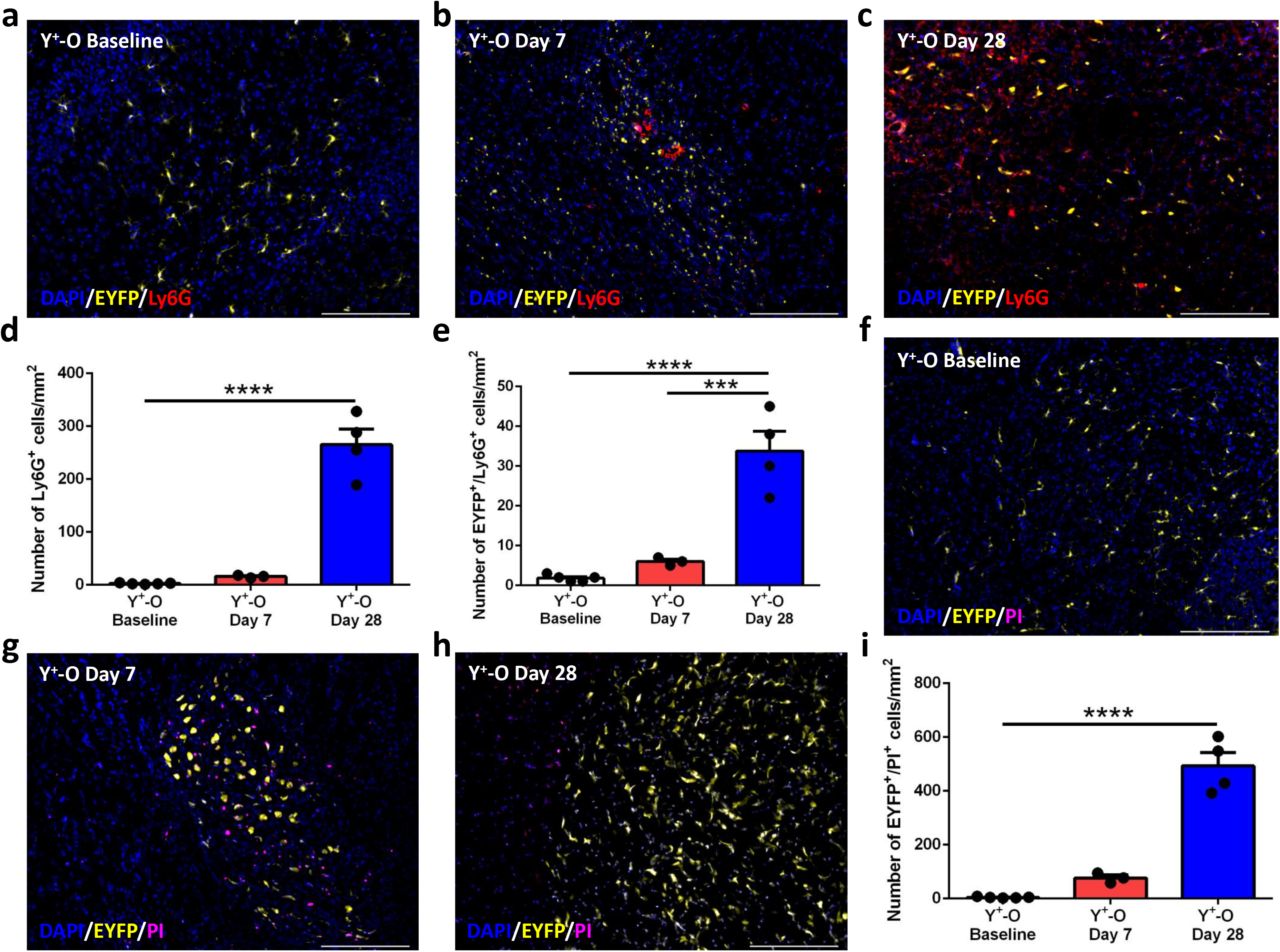
Depletion of YFP^+^ cells by diphtheria toxin is followed by persistence of dead YFP^+^ cells for up to 1 month after initial ablation. (a) Baseline, (b) depletion, and (c) repopulation time points demonstrating YFP^+^ cells and Ly6G^+^ (red) neutrophil numbers in the cerebellum; quantified (d, e). Immunofluorescence staining for cell viability (propidium iodide, purple) during baseline (f), depletion (g), and repopulation (h); quantified (i). Yellow and purple merged will create a white color (nearly all cells). *n* = 5 mice for baseline, *n* = 3 mice for day 7 depletion time point, and *n* = 4 mice for day 28 repopulation time point.

### Young BM-derived microglia-like cells permanently modify hippocampal neurons

Our previous research demonstrated BM reconstitution with young Sca-1^+^ stem cells significantly improved synaptic plasticity and dendritic spine density in pyramidal neurons in aged mice when compared to reconstitution with old Sca-1^+^ stem cells [8]. Since there are only a few cells in the hippocampus 3-months following transplantation, we speculated that the cells likely home to the brain in large numbers during an early time point and influence neuronal function. Indeed, we found a large number of BM-derived YFP^+^ cells in the hippocampus 1-month post-reconstitution (Fig. 7a (left), b)). We were able to deplete these cells from the hippocampus with ∼95% efficiency (Fig. 7b (middle), b) and found the cells begin to repopulate by day 28 (Fig. 7c, (right), b). Next we were curious to see whether depletion and repopulation affected the number of synapses in pyramidal neurons. We found a significantly lower number of dendritic protrusions after depletion (compared to baseline, Fig. 7c), with the number of spines almost recovering back to baseline levels during repopulation (Fig. 7d). Unlike 3-months post-reconstitution, at 1-month post-reconstitution, we found depletion affects novel object recognition (Fig. 7e) and spatial learning and memory (Fig. 7f, g), while repopulation demonstrated partial recovery towards baseline behaviour (Fig. 7e-g).

**Figure 7:**
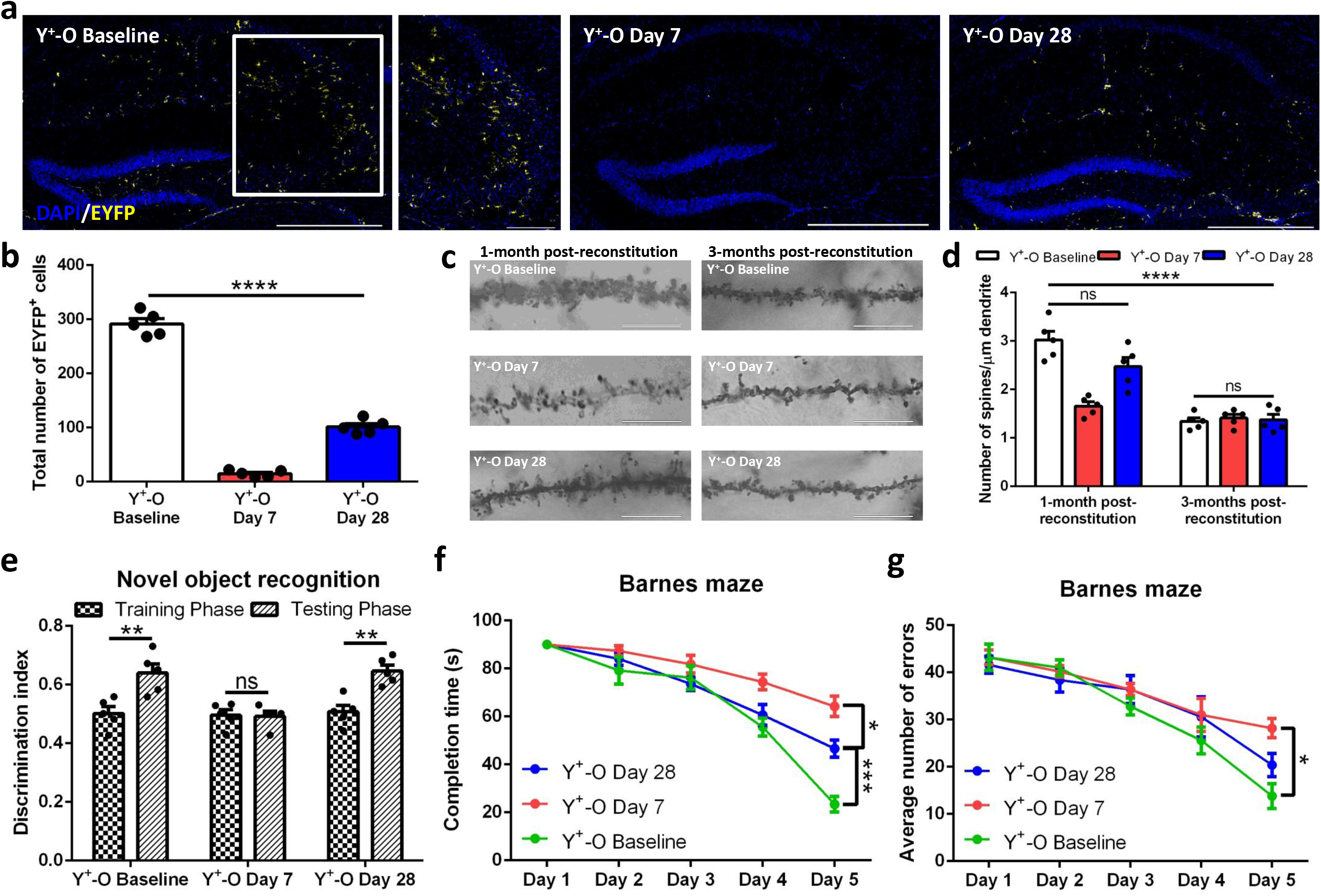
Depletion of BM-derived microglia-like cells 1-month post-reconstitution demonstrates the cells exhibit permanent effects on spine density and cognitive function. (a) Immunofluorescence staining of the hippocampus at baseline 1-month post-reconstitution (left), day 7 depletion (middle), and day 28 repopulation (right), with quantified number of YFP^+^ cells (b). (c) Golgi-Cox staining for dendritic spine densities 1-month and 3-months post-reconstitution with quantified number of spines/µm dendrite (d). Novel object recognition (e) and Barnes maze spatial memory and learning (f, completion time; g, average number of errors) at 1-month post-reconstitution for baseline, depletion, and repopulation time points. Scale bars 200 µm and 20 µm for inset. *n* = 5 mice/group.

### Young BM-derived cells activate genes involved in neurogenesis and spine remodelling

Since we found that hippocampal-dependent learning and memory persists long-term (3-months post-reconstitution) in nearly complete absence of BM-derived cells, we wanted to understand the genetic changes that hippocampal neurons undergo after reconstitution with young Sca-1^+^ stem cells. To study this, we performed a neurogenesis RNA array on the hippocampus of Y^+^-O chimeras 1-month and 3-months post-reconstruction following depletion and at baseline, to represent loss- and gain-of-function, respectively. Our panel revealed young BM-derived cells upregulate genes associated with neurogenesis, neuronal survival and sprouting, and learning and memory (Fig. 8a). Moreover, genes related to synaptic transmission found upregulated TH (Fig. 8b), a mechanism that we proposed previously [8]. Examining a panel of growth factors and cytokines revealed pleiotrophin (PTN, Fig. 8c) and brain-derived neurotrophic factor (BDNF, Fig. 8a). PTN is involved during synaptic plasticity [43], immune cell recruitment [44], and HSC regeneration [45], while BDNF plays a vital role during neurogenesis [12]. Looking at targets related to synaptogenesis and axonogenesis revealed genes involved in neurogenesis and cytoskeleton remodelling of dendritic spines (Fig. 8d). Taken together, these findings demonstrate an early influence (1-month post reconstitution) on the hippocampus by young BM-derived microglia-like cells. This cell influence persists long-term (3-months post-reconstitution) as the genes remained upregulated in the near absence of BM-derived cells from the hippocampus during this time point.

**Figure 8:**
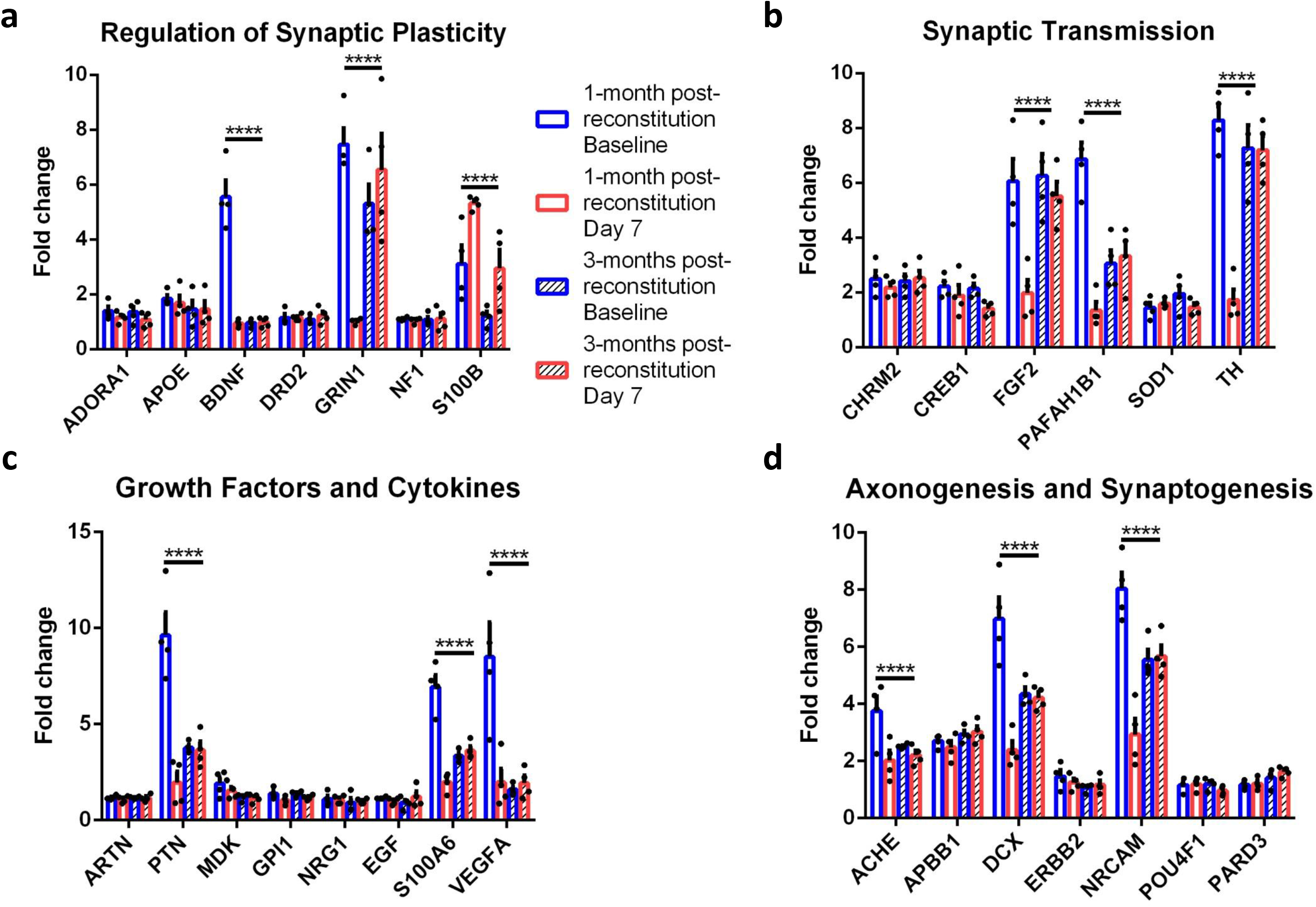
RT-qPCR on hippocampus 1-month and 3-months post-reconstitution in Y^+^-O mice at baseline and after depletion of BM-derived microglia-like cells. Genes grouped according to (a) regulation of synaptic plasticity, (b) synaptic transmission, (c) growth factors and cytokines, and (d) axonogenesis and synaptogenesis for the hippocampus 1-month and 3-months post-reconstitution and after depletion (day 7). *n* = 4 mice/group.

### Young BM-derived microglia-like cells upregulate anti-inflammatory factors

After performing the neurogenesis PCR array, next we were curious to see whether some of the upregulated genes relating to soluble factors were detectable in the hippocampus and whether they were being produced by donor (or host) macrophages. RNA analysis on extracted and purified donor (GFP^+^) macrophages (Fig. 9a) and non-macrophages (Fig. 9b) revealed six different pro- and anti-inflammatory cytokine genes in cells from Y^+^-O and O^+^-O reconstituted brains. Of the six, five of them were anti-inflammatory cytokines/growth factors expressed in young BM-derived microglia-like cells (Fig. 9a), as well as by host microglia (Fig. 9c) from animals reconstituted with young, but not old, Sca-1^+^ cells. In contrast, one of the six detected genes encoded for the pro-inflammatory cytokine, tumor necrosis factor alpha (TNF-α) and was expressed by aged donor microglia-like and non-microglial cells (Fig. 9a, b), as well as by aged host microglia from animals reconstituted with aged Sca-1^+^ cells (Fig. 9c). Of the total of eight factors screened, none of them were significantly upregulated in non-microglial cells from the host (Fig. 9d). Protein screening to confirm gene expression was carried out on ex vivo donor and host microglia and non-microglia, as well as on hippocampal tissue. Here we found that there were several overlapping proteins produced by donor microglia-like cells (Fig. 9e) and donor non-microglia-like cells (Fig. 9f). Based on this data, it would appear that only BDNF was specific to BM-derived microglia-like cells (Fig. 9e) as the remaining factors produced by BM-derived non-microglia can all be produced by monocytes and other immune cells (Fig. 9f). These findings also suggest that young BM-derived microglia-like cells might confer expression of these proteins in host microglia (Fig. 9g), a phenomenon that has been previously mentioned [46], but not host non-microglia (Fig. 9h). Lastly, the proteins produced by microglia-like cells are also detected in the hippocampus 1-month post-reconstitution, but not so much after depletion (Fig. 9i).

**Figure 9:**
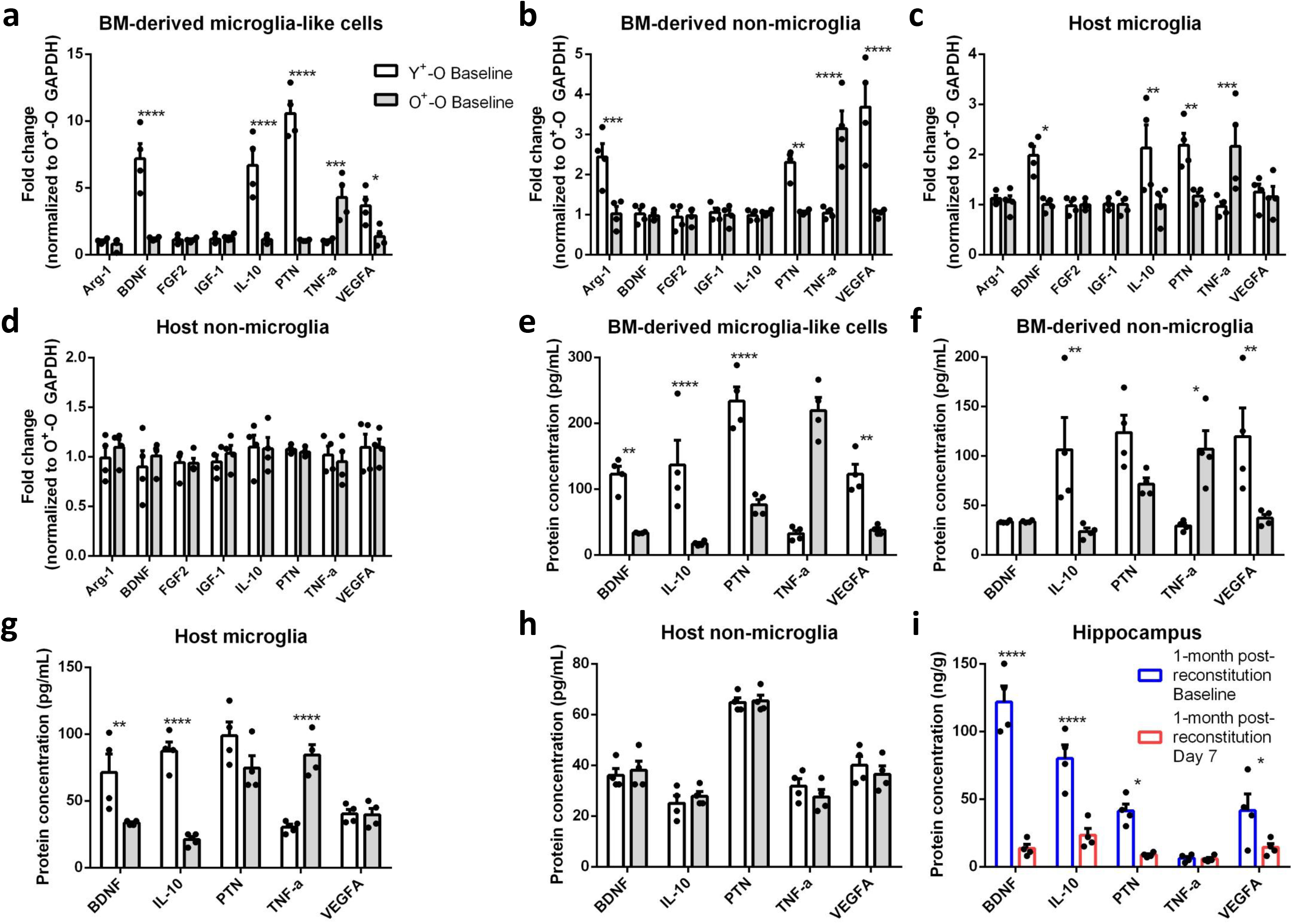
RT-qPCR and protein analysis on cells and tissue from the brain for inflammatory cytokines and growth factors. (a) RT-qPCR on BM-derived microglia-like cells, (b) BM-derived non-microglia, (c), host microglia, and (d) host non-microglia for eight pro- and anti-inflammatory cytokines and growth factors. (e) Verification of RNA expression using protein analysis for BM-derived microglia-like cells, (f) BM-derived non-microglia, (g), host microglia, and (h) host non-microglia for eight pro- and anti-inflammatory cytokines and growth factors. (i) Confirmation of protein expression in the hippocampus 1-month post-reconstitution at baseline and during the day 7 depletion time point. *n* = 4 mice/group.

### Young BM-derived microglia-like cells promote dendritogenesis via the BDNF-TrkB pathway

We demonstrated that the presence of young BM-derived microglia-like cells is vital during an early time point (1-month post-reconstitution) in order to confer permanent changes to neurons due to the large presence of these donor cells not being required long-term (3-months post-reconstruction) for maintenance of improved cognitive function and synaptic plasticity. By which mean(s) this occurs has yet to be determined. Since we found that BDNF is the only unique factor produced by young donor BM-derived cells, we decided to block the binding of this growth factor to its receptor, tropomyosin-related kinase B (TrkB), using the antagonist ANA-12. This antagonist was previously reported to block said receptors with high potency and specificity, while affecting cognitive behavior without compromising neuronal survivability [47]. To achieve this, ANA-12 was administered every day for five consecutive days and behavior was carried out (Fig. 10a). Once behavioral testing concluded, the animals were sacrificed and brains were collected for dendritic spine analysis using Golgi-Cox staining (Fig. 10b, overview; Fig. 10c, hippocampus (top)). We found that blocking TrkB receptors using ANA-12 inhibited dendritogenesis 1-month post-reconstitution when compared to saline-injected control in Y^+^-O mice (Fig. 10c (middle, bottom), d). Moreover, we found that behavioral performance was significantly compromised in ANA-12-injected Y^+^-O mice on hippocampal-dependent novel object recognition (Fig. 10e) and Barnes maze spatial memory and learning for average number of errors (Fig. 10f) and maze completion time (Fig. 10g). Taken together, these findings reveal at least one mechanism of action by which young BM-derived cells benefit neuronal and cognitive function in the aged brain: by migrating to the hippocampus early after reconstitution and secreting BDNF that is required for promoting long-term dendritogenesis.

**Figure 10:**
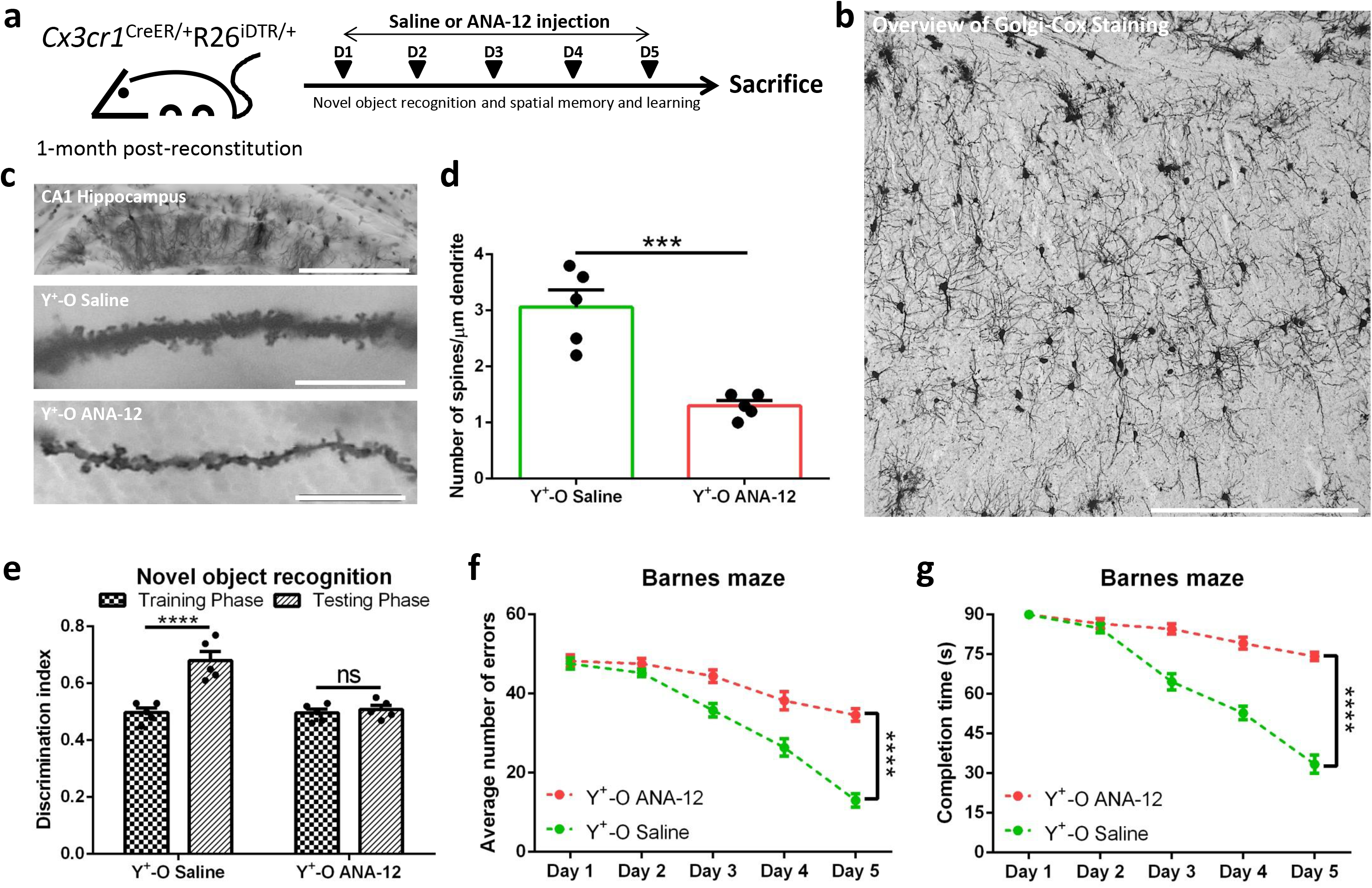
Blocking BDNF receptor binding with ANA-12 in the hippocampus 1-month post-reconstitution inhibits dendritogenesis and cognitive improvements that are driven by young BM-derived microglia-like cells. (a) Cartoon representation of timeline regarding drug administration and behavioral testing. (b) Brain overview of Golgi-Cox staining with example of pyramidal neuron staining in the CA1 hippocampus (c, top). Dendritic spine analysis in Y^+^-O saline-treated group (c, middle) and Y^+^-O ANA-12 injected group (c, bottom), quantified (d). Novel object recognition (e), Barnes maze average number of errors and (g) completion time for Y^+^-O saline-treated group and Y^+^-O ANA-12 injected group. *n* = 5 mice/group.

## Discussion

In this study we demonstrated that partial depletion of young BM-derived microglial-like cells from aged recipients does not reverse hippocampal-driven behavioral performance to levels present in aged recipient controls of old donor cells at 3-months post-reconstitution, but not at 1-month post-reconstitution. This depletion at 3-months is accompanied by a severe global inflammatory response that seems to primarily be mediated by neutrophils. This appears to be influenced by the age of the animal as past microglia depletion data on young reconstituted mice show that neutrophil numbers in the brain fail to significantly increase above baseline levels following depletion [38]. Moreover, contradictory to previous microglia/macrophage depletion studies, we were unable to see repopulation of the macrophage niche in the aged brain after depletion and 3-months post-reconstitution; neither from brain-resident nor peripherally-derived cells. Our data shows that ablation of cells from the aged brain is not compensated by an influx of monocytes that can potentially contribute to the repopulation of the CX3CR1 compartment.

Depletion of macrophages from the brain using tamoxifen-induced expression of diphtheria toxin receptors has existed for nearly a decade [12]. Although we achieved >50% depletion of BM-derived macrophages 3-months post-reconstruction, other studies have been able to reach >90% depletion during this time point. This difference in depletion efficiency could be explained by a shift towards using the *R26*^DTA^ [38] model of depletion, rather than the *R26*^DTR^. One of the limitations of this study is that we used the latter model, of which the former results in self-depletion of targeted cells via the production of intracellular diphtheria toxin following exposure to tamoxifen – eliminating the need for exogenous application of diphtheria toxin. Despite using a model of smaller efficiency of depletion, we hypothesize that this may not have made a difference in overall percent cell depletion as our data nonetheless demonstrated impaired clearance of dead YFP^+^ cells.

In this study we discovered that age of the animal where cells are being depleted, as well as the time of depletion, is a vital variable in determining depletion efficiency and subsequent ability to repopulate the macrophage niche. Past studies have failed to take the age of the animal into consideration [18]. Normally after depletion, microglia begin to rapidly repopulate by 48-72 hours from remaining self-proliferating microglia that did not get depleted and from peripherally-derived precursor cells [42]. The rapid expansion and repopulation of the microglial niche following depletion has been one of the major limitations regarding the depletion of these cells in order to study the effects their absence has on animal and cell physiology. Here we report a novel and crucial finding demonstrating that the macrophage/microglial niche fails to repopulate following depletion 3-months post-reconstitution in aged animals. Neither blood-borne nor peripheral sources are able to contribute towards rebuilding the CX3CR1 compartment after ablation. Although there may be several reasons for this, our data demonstrates the persistence of BM-derived microglial-like cells following diphtheria toxin-induced cell death. As such, repopulation of the niche may not be made possible by a lack of physical availability for newly formed and migrated cells. Moreover, extensive inflammation during the repopulation time point is met with a cell-unfriendly environment. Interestingly, even after phagocytosis of YFP^+^ cells by neutrophils, we find the persistence of YFP signal in these cells, further demonstrating the challenges the aged immune system faces in response to global cell depletion and the demand to repopulate this niche.

Lastly, selective depletion and drug antagonist data 1-month post-reconstitution demonstrated a vital time point in which BM-derived microglia-like cells induce dendritogenesis in hippocampal neurons and enhancement of cognitive function via the BDNF-TrkB signalling pathway. These early-acquired beneficial effects persist long-term even in the near-absence of donor cells in the hippocampus, demonstrating one mechanism by which a single intravenous injection of Sca-1^+^ hematopoietic stem cells greatly benefits cognitive function restoration in aged recipients.

## Acknowledgments

We thank Gregory Wlodarek for programming custom Python scripts to analyze and quantify animal behaviour.

## Author contributions

L.W. designed the experiments, carried out and analyzed biochemical and behavioural paradigms, and wrote the manuscript. R.K.L contributed to study design, data interpretation and preparation of the manuscript.

## Conflicts of interest

The authors have no conflicts of interest to declare.

## Data availability statement

All data are available upon reasonable request from the principal investigator.

## Grant acknowledgement

This work was supported by grants from the Canadian Institutes of Health Research (332652) to R.K.L.

## Notes

### Competing Interest Statement

The authors have declared no competing interest.

